# Peptide-MHC Structure Prediction With Mixed Residue and Atom Graph Neural Network

**DOI:** 10.1101/2022.11.23.517618

**Authors:** Antoine P. Delaunay, Yunguan Fu, Alberto Bégué, Robert McHardy, Bachir A. Djermani, Michael Rooney, Andrey Tovchigrechko, Liviu Copoiu, Marcin J. Skwark, Nicolas Lopez Carranza, Maren Lang, Karim Beguir, Uğur Şahin

## Abstract

Neoantigen-targeting vaccines have achieved breakthrough success in cancer immunotherapy by eliciting immune responses against neoantigens, which are proteins uniquely produced by cancer cells. During the immune response, the interactions between peptides and major histocompatibility complexes (MHC) play an important role as peptides must be bound and presented by MHC to be recognised by the immune system. However, only limited experimentally determined peptide-MHC (pMHC) structures are available, and *in-silico* structure modelling is therefore used for studying their interactions. Current approaches mainly use Monte Carlo sampling and energy minimisation, and are often computationally expensive. On the other hand, the advent of large high-quality proteomic data sets has led to an unprecedented opportunity for deep learning-based methods with pMHC structure prediction becoming feasible with these trained protein folding models. In this work, we present a graph neural network-based model for pMHC structure prediction, which takes an amino acid-level pMHC graph and an atomic-level peptide graph as inputs and predicts the peptide backbone conformation. With a novel weighted reconstruction loss, the trained model achieved a similar accuracy to AlphaFold 2, requiring only 1.7M learnable parameters compared to 93M, representing a more than 98% reduction in the number of required parameters.

## 1 Introduction

Cancer immunotherapeutics have revolutionised the field of oncology by using the patient’s immune system to trigger tumour regression [Finck et al., 2020, Waldman et al., 2020]. Among them, cancer vaccines aim to induce and enhance tumour-specific T-cell responses in patients by delivering targeted immunogenic neoepitopes [Blass and Ott, 2021]. The cellular immune response activation towards an antigen relies on a series of tightly regulated biological processes whose keystones are the major histocompatibility complexes (MHC). These essential cell surface proteins bind peptides and display them to the intercellular space where they can interact with immune cells. The features underlying the binding and presentation of antigenic peptides are therefore key to understanding the immune response and have valuable health implications.

However, the MHC is a highly polymorphic protein and there are 20^9^ possible 9-mer peptide sequences, with limited structural information available in public databases [Berman et al., 2000]. In-silico structure modelling methods are thereby applied for predicting peptide-MHC (pMHC) structures. State-of-the-art methods mostly rely on energy minimisation strategies via random sampling but require a long computation time, making scalability a challenge [Abella et al., 2019, Parizi et al., 2022]. The advent of large high-quality proteomic data has led to an unprecedented opportunity for deep learning-based protein folding methods such as AlphaFold 2 [Jumper et al., 2021], OmegaFold [Wu et al., 2022], and ESMFold [Lin et al., 2022], which have been proven to predict reliable and accurate protein structures. Furthermore, these trained models can be fine-tuned on pMHC structures for other downstream tasks such as binding prediction [Motmaen et al., 2022].

In this paper, we propose a graph neural network (GNN) that takes an amino acid residue graph for the pMHC interface and an atom graph for the peptide as inputs to predict the peptide backbone conformation. A post-processing step recovers the full-atom coordinates and constructs the entire pMHC structure. By training on a novel weighted reconstruction loss, the proposed GNN predicted accurate structures with similar performance compared to AlphaFold 2 [Jumper et al., 2021] and pMHC fine-tuned AlphaFold 2 [Motmaen et al., 2022] with only 1.7M learnable parameters, which represents a more than 98% reduction in the number of parameters. We demonstrate that the predicted structures are closer to native structures, both in terms of geometry and biological consistency. Furthermore, we show that our method considerably improves upon others in terms of fulfilling biological constraints on peptide positioning within certain areas of the MHC called binding pockets [Nguyen et al., 2021].

## 2 Methods

### 2.1 pMHC Graph Representation

Unlike protein folding models that predict structures from amino acid sequences and pairwise features of template structures, in this work, we propose to input an initial conformation by replacing the residues in the template structure with target ones. Specifically, given a target pMHC, we identify a template pMHC structure from a set of experimentally determined pMHC structures via peptide sequence similarity and replace the amino acid residues with target ones for both peptide and MHC. In other words, we input the target pMHC sequence with the conformation from the selected template.

The resulting pMHC structure is represented by two graphs, one representing the pMHC interface at the amino acid-level to capture the interaction between the peptide and MHC, and another one representing the peptide on the atom-level to refine the information for conformation prediction.

The pMHC graph consists of the amino acid residues of the peptide together with the *α*_1_ and *α*_2_ domains of the MHC [Wilson and Fremont, 1993]. Nodes represent residues, with edges connecting nodes if the distance between the *C*_*α*_ of the corresponding residues is smaller than 8 Å, following Xia and Ku [2021]. For each node, the features include the type of amino acid, Kidera factors [Kidera et al., 1985], Atchley factors [Atchley et al., 2005], and the coordinates of the *C*_*α*_, *O, C*_carbo_, *N* and *C*_*β*_ atoms of the residue. For glycine, a virtual *C*_*β*_ position is computed following Cock et al. [2009]. For each edge, the features include the edge classes depending on the bond type (covalent or non-covalent) and the chains (peptide or MHC) of the nodes it connects to.

The peptide graph consists of the backbone atoms (*C*_*α*_, *O, C*_carbo_, and *N*) and *C*_*β*_ atoms. Nodes represent atoms, with edges connecting nodes if a covalent bond exists between the corresponding atoms. The node features encode the atom classes and the edge features encode the edge classes depending on whether it is a single or double covalent bond and whether the atoms it connects come from the same residue.

### 2.2 Neural Network Architecture

The neural network consists of three stages (Figure 1). First, the amino acid-level pMHC graph is fed into a GNN block and each amino acid residue is represented by a node embedding. Meanwhile the atom-level peptide graph is constructed where atom features are encoded and summed with their corresponding residue embeddings. The resulting embeddings together with the atom-level peptide graph are fed into another GNN block to refine the atom embeddings. Finally, each atom’s coordinates are predicted from their embeddings via two shared dense layers.

**Figure 1:**
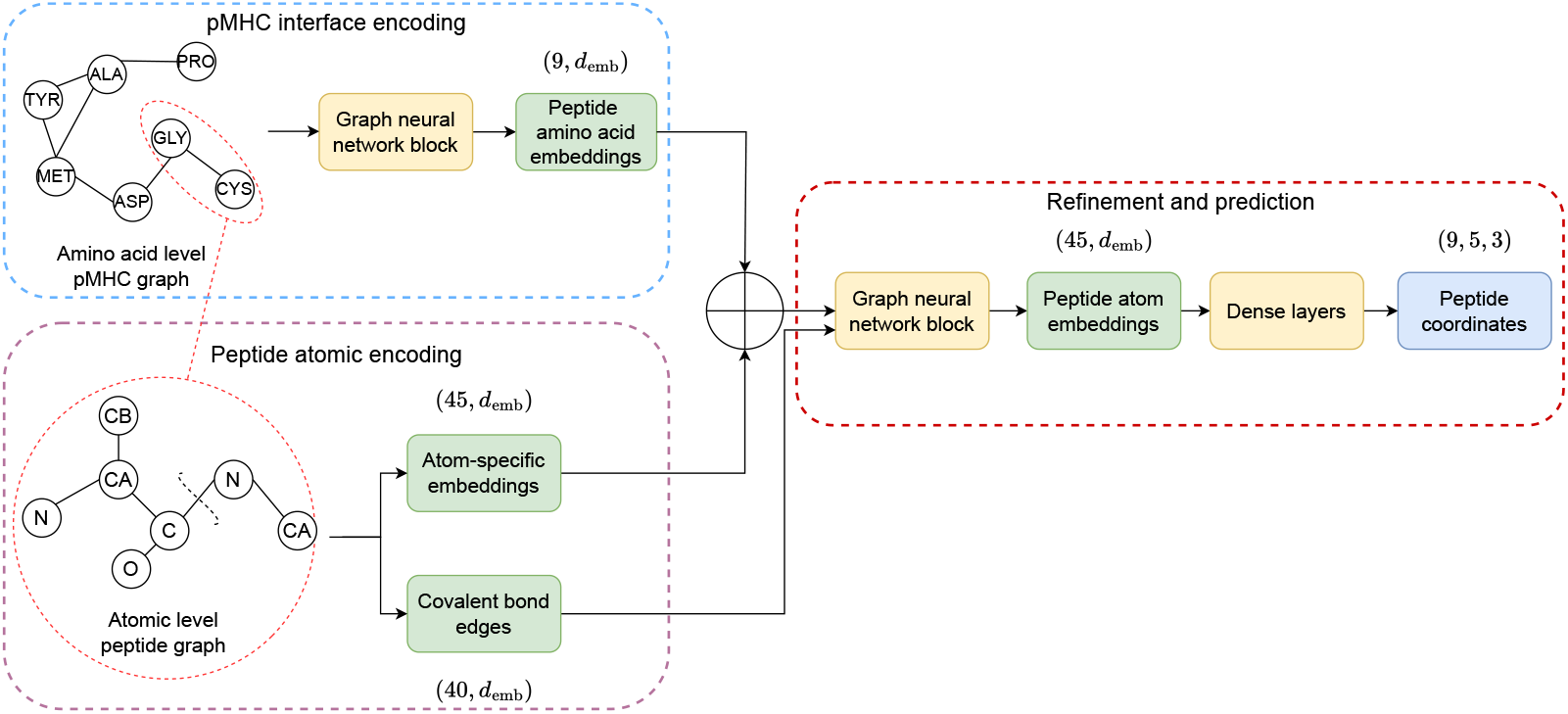
Neural Network Architecture. Each model stage (pMHC interface encoding, peptide atomic encoding, refinement and prediction) are represented by a rectangular dashed box in light blue, mauve, and red, respectively. Model layers and layers’ inputs/outputs are represented by rectangular colour-filled boxes, in yellow and green, respectively. The blue-filled rectangular box represents the final model output (peptide atom coordinates).

The key component is the GNN block which consists of four message passing layers with an multi-head attention mechanism [Shi et al., 2021]. The message passing layer is wrapped into a Transformer-like architecture involving a ReLU-activated feed-forward network. To reduce the impact of distant neighbours, the distance between each pair of nodes is considered in the denominator inside the message passing layer:

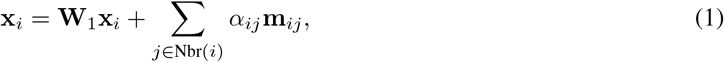

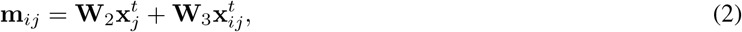

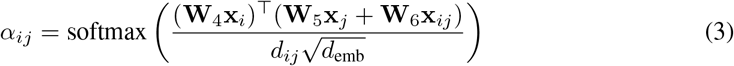

where **W**_1_, **W**_2_, **W**_3_, **W**_4_, **W**_5_ and **W**_6_ are learnable parameters of shape *d*_emb_ × *d*_emb_; **x**_*i*_ is the embedding for node *i*; **x**_*ij*_ is the embedding for edge between node *i* and *j*, and *j* iterates through all neighbour nodes of *i*; **m**_*ij*_, *α*_*ij*_, and *d*_*ij*_ are the message, attention, distance from node *j* to *i*, respectively; and *d*_emb_ is the dimension of the embedding.

### 2.3 Loss

We propose a novel loss *L*_Struct_ to ensure chemical and geometric constraints on the peptide structure. It includes 1) *L*_MSE_ (Appendix A.1), a mean squared error (MSE) of atom coordinates between ground truth and prediction, 2) *L*_Intra_ (Appendix A.2), a MSE of pairwise distances between atoms inside each residue, 3) *L*_Inter_ (Appendix A.3), a MSE of pairwise distances between atoms across different residues, 4) *L*_Dihedral_ (Appendix A.4), a MSE of the trigonometric functions (cos and sin) of dihedral angles:

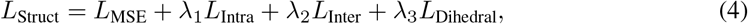

where *λ*_1_, *λ*_2_ and *λ*_3_ are hyper-parameters.

### 2.4 Post-processing

For each peptide residue, as the network outputs only the backbone and *C*_*β*_ atom coordinates, calculation of side chain atoms is necessary to obtain a full-atomic structure. We use the Rosetta Packer algorithm [Leaver-Fay et al., 2011] which selects the side chain conformations from the 2010 BBDep Rotamer library [Shapovalov and Dunbrack, 2011], such that the overall energy (Rosetta REF15 energy [Alford et al., 2017]) of the structure is minimised, similar to the post-processing applied by Jumper et al. [2021] in AlphaFold 2. During the post-processing, only side chain atoms are updated, the predicted backbone atoms remain unchanged.

## 3 Experiment Setting

### 3.1 Data

We focus on MHC class I with a data set of 749 pMHC crystal structures retrieved from the Research Collaboratory for Structural Bioinformatics (RCSB) Protein Data Bank (PDB) [Berman et al., 2000]. Structures with any missing backbone or *C*_*β*_ atom and with non-9-mer peptides were removed. The *β*-2 microglobulin chain was not used in this work. After this process, 458 structures were kept. The structures were further split into train, validation, and test sets such that the test set matched the same as the one used by Motmaen et al. [2022] with no peptide shared between the splits. The resulting data set contains 293, 36, and 38 structures for train, validation, and test split, respectively. PDB IDs and amino acid frequencies of each subset are provided in Appendix B. All structures were aligned into the same coordinate system based only on the MHC *C*_*α*_ atoms with respect to a randomly chosen structure (PDB code: 1AKJ).

### 3.2 Metrics

We calculated the root mean square deviation (RMSD) on backbone and full-atom peptide structures for evaluation. Structures predicted by AlphaFold 2 and OmegaFold were aligned with respect to the 1AKJ MHC chain prior to evaluation. Additionally, we considered a mean average error (MAE) of the dihedral angles *ϕ* and *ψ*, averaged across the peptide. As the peptide positioning has to fulfil biological constraints and be inserted into MHC surrounding areas called binding pockets [Nguyen et al., 2021], we evaluated whether the predicted structures exhibit accurate binding positions by calculating the MAE on distances from the peptide to the MHC binding pockets (Appendix C). The total reweighted and attractive energy scores with Rosetta [Raveh et al., 2011] were reported to quantitatively assess the biological consistency of the structures.

### 3.3 Implementation

The GNN model was implemented with PyTorch Geometric [Fey and Lenssen, 2019] and trained on a Nvidia A100 40GB GPU. The optimiser was Adam with an initial learning rate of 3 × 10^−4^. Hyper-parameters were empirically set without extensive tuning (Appendix D). The official implementation of AlphaFold 2 monomer [Jumper et al., 2021], pMHC fine-tuned AlphaFold 2 [Motmaen et al., 2022], and OmegaFold [Wu et al., 2022] were used for benchmarking.

## 4 Results

The metrics of the proposed model together with the baselines AlphaFold 2 [Jumper et al., 2021], pMHC fine-tuned AlphaFold 2 [Motmaen et al., 2022], and OmegaFold [Wu et al., 2022] are reported in Table 1 and their distributions in Figure 6 (Appendix E). For the GNN, results for the first seed are given in Table 1 and for four other seeds in Table 9 (Appendix E). The proposed method achieved on average 1.26 Å and 2.51 Å backbone and full-atom RMSD, respectively. This is comparable with the AlphaFold 2-based counterparts, despite having only 1.7M learnable parameters compared to 93M in AlphaFold 2. High backbone accuracy is necessary since full-atom structures are produced based on the backbone coordinates, therefore small errors in the backbone might propagate and have large effects on the side-chains. OmegaFold, on the other hand, generated inaccurate results with an RMSD of >10 Å (Figure 7 in Appendix E). The average dihedral MAE of the GNN is 26°, which is acceptable since the Packer algorithm uses a dihedral angle resolution of 10°. When comparing the distance to binding pockets, the proposed method achieved lower MAE, meaning that our method predicted structures better satisfying the constraints on the anchor residues. In terms of energies, all models predicted structures with higher energy scores than native, but the proposed method achieved the smallest difference, suggesting that structures predicted by the GNN are energetically more consistent with native structures than other methods. Examples of the predicted pMHC structures are visualised in Figure 2. Ablation studies were performed to analyse the impact of the proposed novel loss functions. The results are summarised in Table 10 in Appendix F, showing that additional loss terms led to improved structure geometry (dihedral angles) and more energetically consistent structures. Binding pockets were less impacted by the new loss terms, with the overall peptide position in the binding groove predominantly determined via the MSE loss term.

**Table 1:**
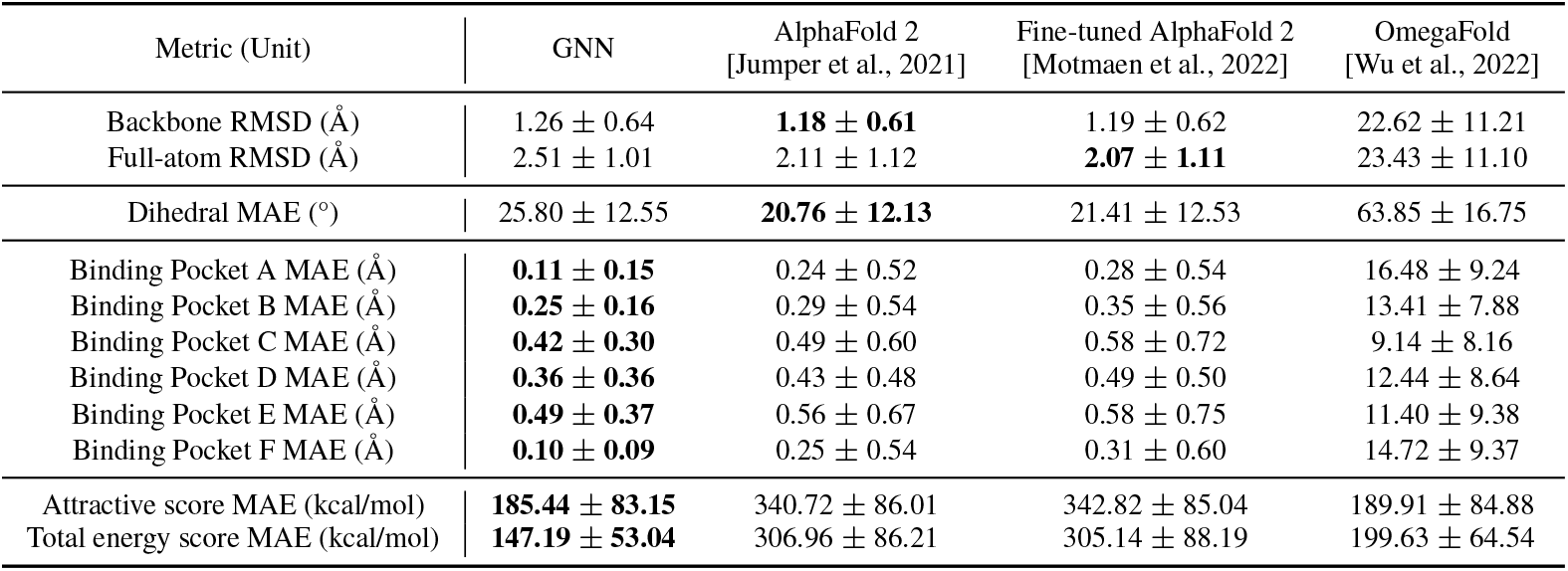
Metrics on test set per model. The reported values are average with standard deviation.

**Figure 2:**
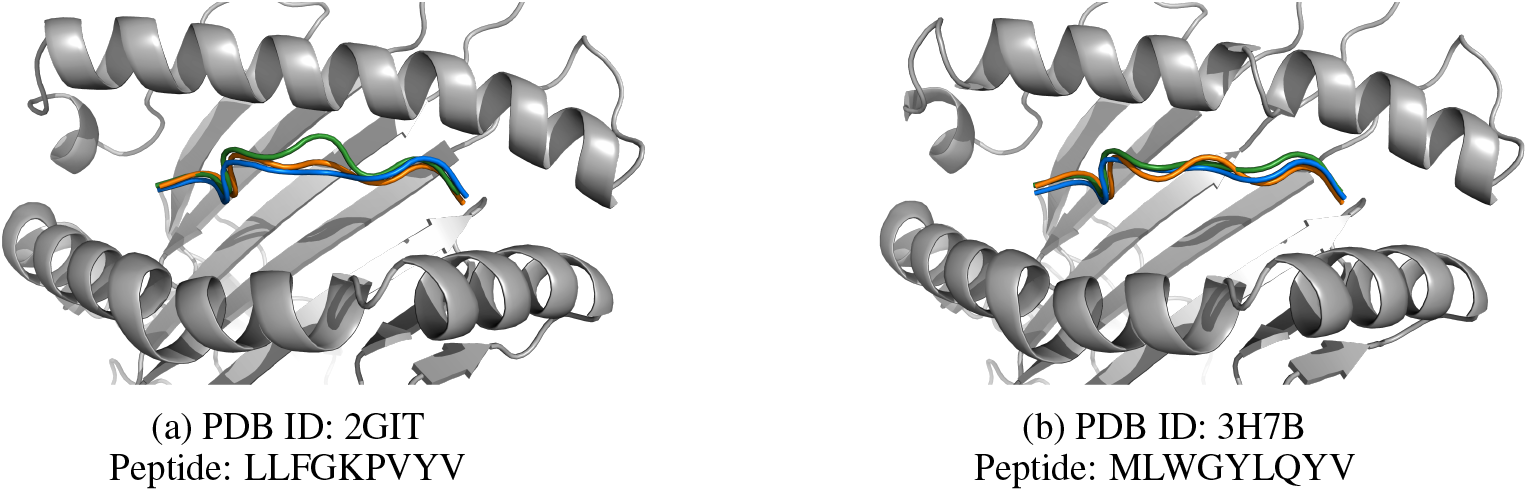
Examples of predicted structures. Experimental, GNN, and AlphaFold 2 fine-tuned predicted peptide are represented in green, blue, and orange, respectively. The MHC chain is represented in grey. AlphaFold 2 structures are not represented to simplify visualisation. GNN matches AlphaFold 2 fine-tuned structures and captures most of the peptide backbone shape and curvatures.

## 5 Conclusion and Discussion

In this work, we proposed a novel graph neural network architecture taking both residue-level and atom-level graphs to predict full-atomic pMHC structures. By minimising the composed losses on atom coordinates, pairwise distances, and dihedral angles, we demonstrated that a lightweight model with only 1.7M parameters is capable of producing accurate and biologically consistent structures, which is comparable to large protein folding models such as AlphaFold 2 (93M parameters). While the proposed method achieved plausible results, it solely predicts the backbone and *C*_*β*_ atoms for the peptide and relies on the Rosetta Packer algorithm to obtain full-atomic structures. Extending the network to predict full-atom pMHC structures may enable us to achieve higher accuracy, and to further reduce inference time. Moreover, the experiments were focused only on the interactions between 9-mer peptides and MHC class I. Future work could include extending the model to peptides of variable length and MHC class II to build a universal structure prediction model for pMHC complexes.

## A Loss

### A.1 Mean Squared Error Loss

The role of the mean squared error loss is to give the correct overall shape and position in the space of the peptide atoms. There are 9 amino acids and 5 atoms per amino acid. Let 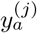 and 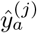 denote the true and predicted coordinates of the *j*^*th*^ atom in the *a*^*th*^ amino acid of the peptide, respectively. As some peptide residues are more variable than others, residue-wise weights (denoted by *w*_*a*_) are added to strengthen the attention given to the middle amino acids. Raw residue-wise weights are calculated by running the model with uniform weighting (i.e. *w*_*a*_ = 1*/*9) and then extracting the residue-wise MSE loss on the validation subset. Weights *w*_*a*_ are normalised to sum to 1 and the values are provided in Table 2. The formula of this loss is given by:

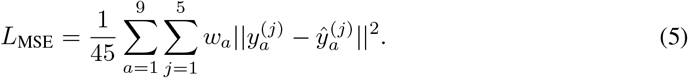

**Table 2:**
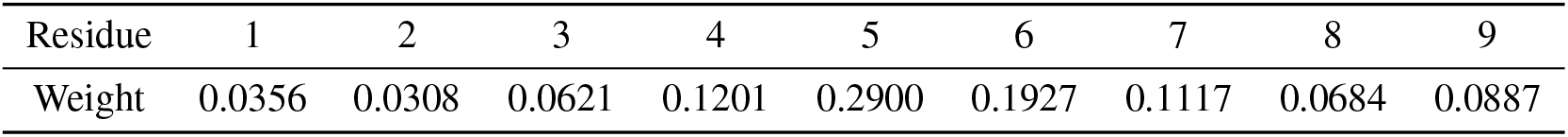
MSE weights values per amino acid.

| Residue | 1 | 2 | 3 | 4 | 5 | 6 | 7 | 8 | 9 |
| --- | --- | --- | --- | --- | --- | --- | --- | --- | --- |
| Weight | 0.0356 | 0.0308 | 0.0621 | 0.1201 | 0.2900 | 0.1927 | 0.1117 | 0.0684 | 0.0887 |

### A.2 Intra-residue Loss

Let 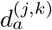 and 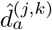 denote the distances between the *j*^*th*^ and *k*^*th*^ atoms in the *a*^*th*^ amino acid of the peptide for the ground truth and predicted structures, respectively:

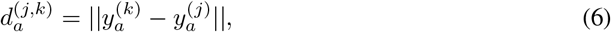

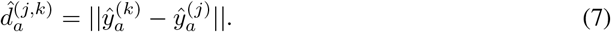

The intra-residue loss is defined as (Figure 3):

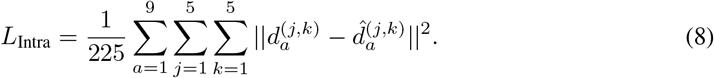

**Figure 3:**
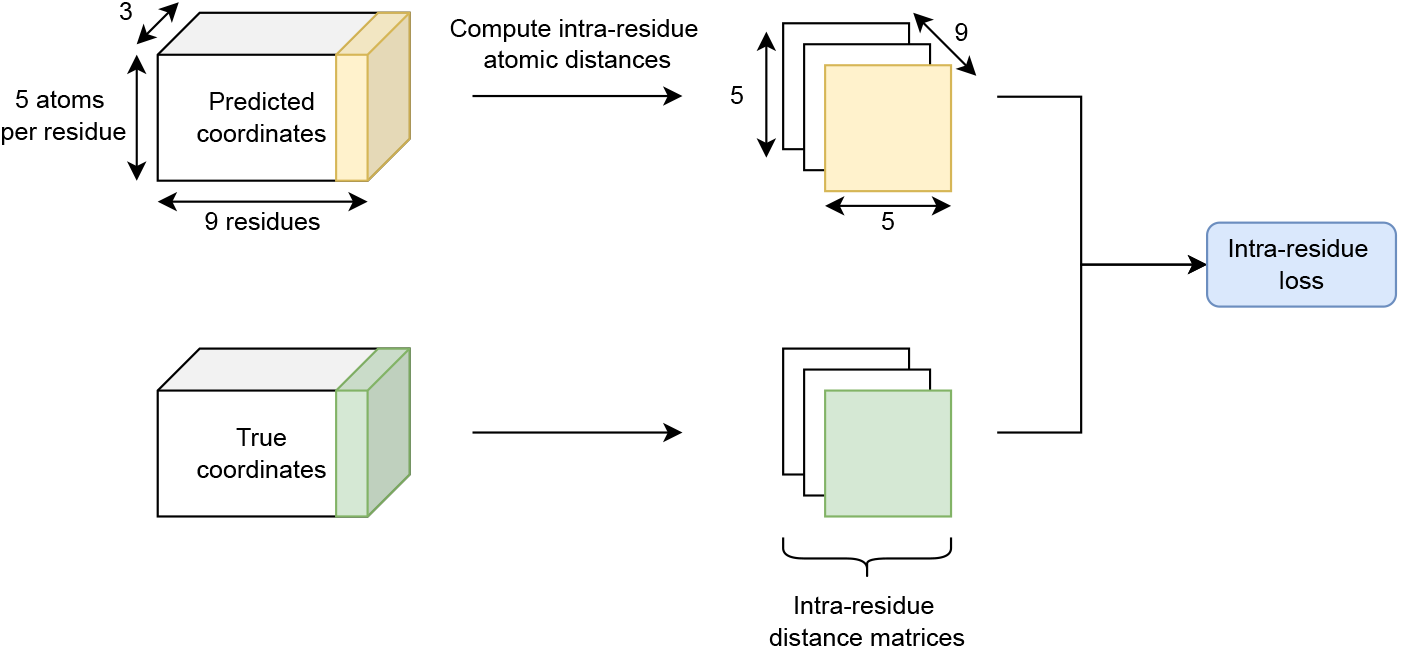
Intra-residue loss computation process

The intra-residue loss adds constraints on the local geometry inside each peptide amino acid residue.

### A.3 Inter-residue Loss

Let *d*_*a,b*_ and 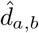 denote the distances between the *C*_*α*_ atoms of the *a*^*th*^ and *b*^*th*^ amino acids of the peptide for the ground truth and predicted structures, respectively:

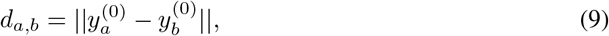

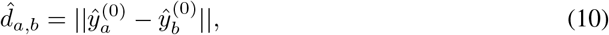

where 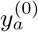 and 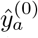 represent the coordinates of the *C*^*α*^ atom for *a*^*th*^ amino acid of peptide for the ground truth and predicted structures, respectively. The inter-residue loss is defined as (Figure 4):

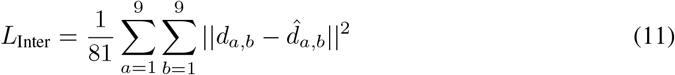

**Figure 4:**
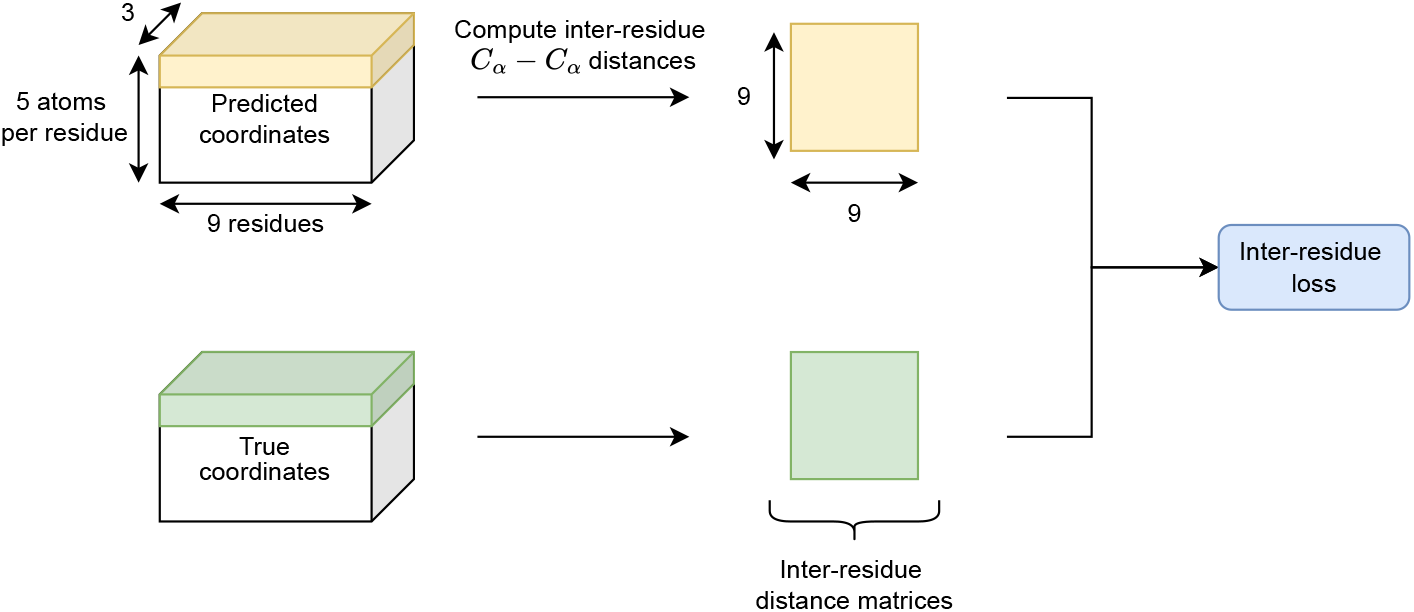
Inter-residue loss computation process

The inter-residue loss adds constraints on the geometry between peptide amino acid residues.

### A.4 Dihedral Loss

Let *ϕ*^*j*^ and *ψ*^*j*^ denote the *j*^*th*^ dihedral angles of rotation over the *N* − *C*_*α*_ and *C*_*α*_ − *C* covalent bonds of the peptide [Ramachandran et al., 1963]. Following Xia and Ku [2021], the dihedral loss is defined as:

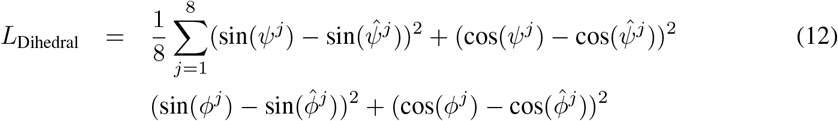

## B Data

The PDB IDs for the training, validation, and test sets are listed in Table 3, Table 4, and Table 5, respectively. The amino acid frequencies at each peptide position are illustrated in Figure 5 for each data subset.

**Table 3:**
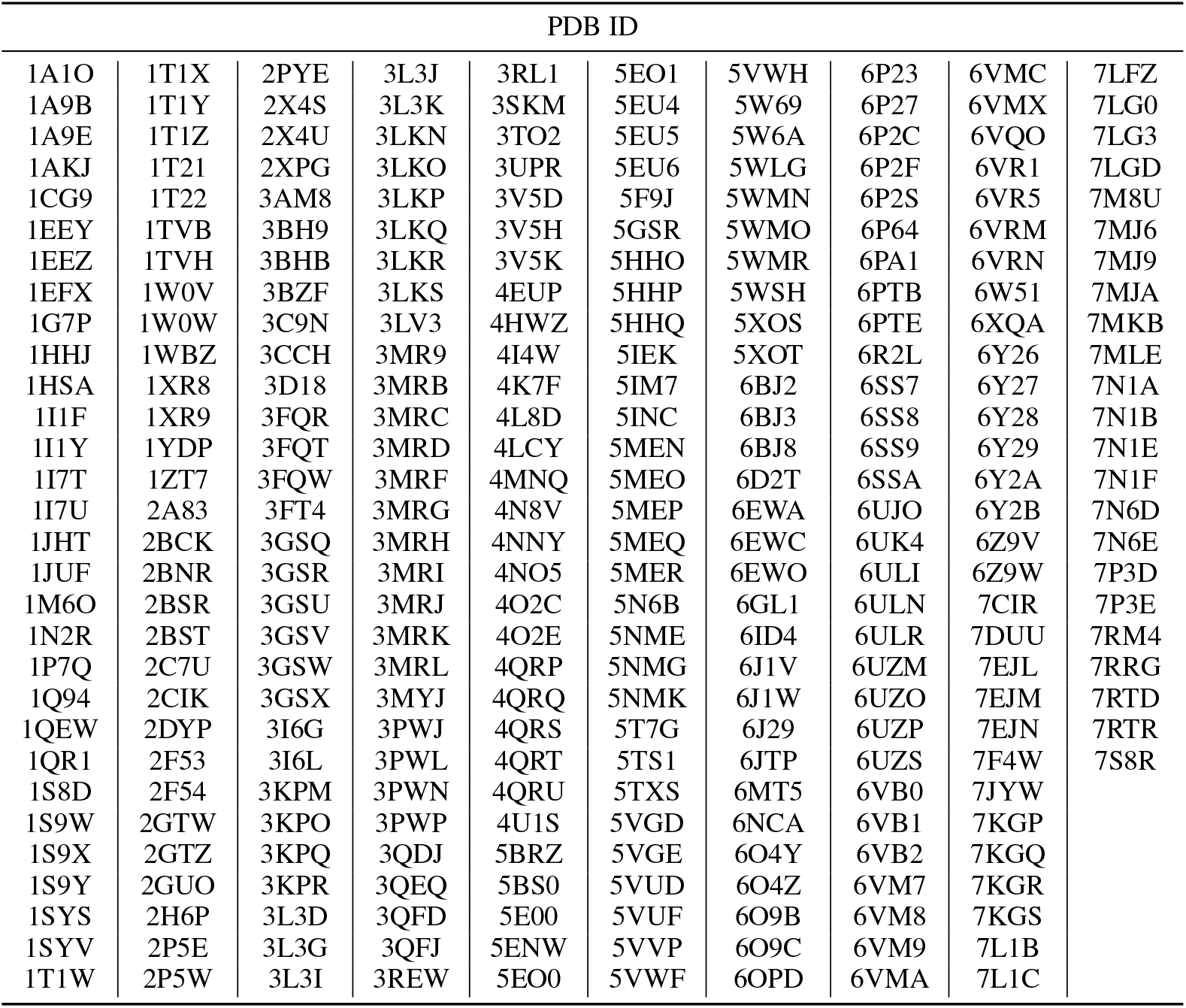
Train set PDB IDs.

**Table 4:**
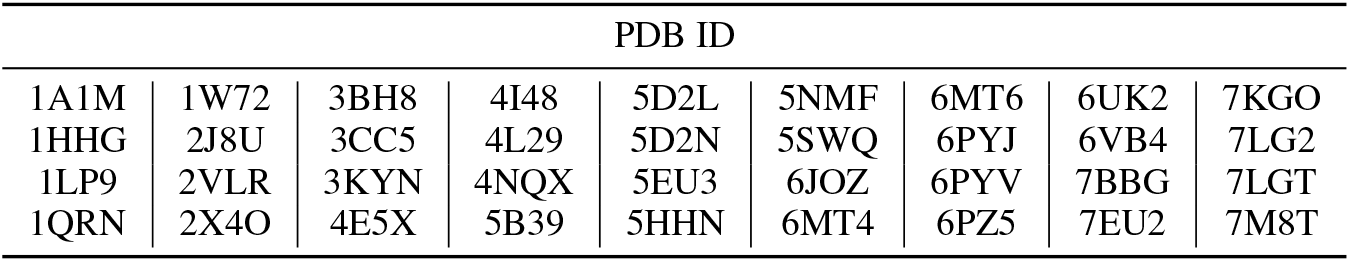
Validation set PDB IDs.

| PDB ID |  |  |  |  |  |  |  |  |
| --- | --- | --- | --- | --- | --- | --- | --- | --- |
| 1A1M | 1W72 | 3BH8 | 4I48 | 5D2L | 5NMF | 6MT6 | 6UK2 | 7KGO |
| 1HHG | 2J8U | 3CC5 | 4L29 | 5D2N | 5SWQ | 6PYJ | 6VB4 | 7LG2 |
| 1LP9 | 2VLR | 3KYN | 4NQX | 5EU3 | 6JOZ | 6PYV | 7BBG | 7LGT |
| 1QRN | 2X4O | 4E5X | 5B39 | 5HHN | 6MT4 | 6PZ5 | 7EU2 | 7M8T |

**Table 5:** Test set PDB IDs.

| PDB ID |  |  |  |  |  |  |  |  |  |
| --- | --- | --- | --- | --- | --- | --- | --- | --- | --- |
| 1E27 | 1QQD | 2FWO | 3D25 | 3KYO | 4QRR | 5DEG | 5TRZ | 6G9Q | 6VB7 |
| 1I7R | 1VGK | 2GIT | 3H7B | 3MRE | 4U1H | 5FA3 | 5U98 | 6GH1 | 7JYV |
| 1JGE | 1X7Q | 3BVN | 3KLA | 3SKO | 4U1N | 5IB2 | 5VWJ | 6J2A |  |
| 1JPG | 2BVP | 3BXN | 3KPP | 4HX1 | 4Z77 | 5IND | 5WMQ | 6Q3S |  |

**Table 6:** Binding pocket peptide and MHC anchor residues from Nguyen et al. [2021].

| Binding pocket | Peptide residue(s) | MHC anchor residues |
| --- | --- | --- |
| A | 1 | 5, 7, 59, 63, 66, 159, 163, 167, 171 |
| B | 2 | 7, 9, 24, 34, 45, 63, 66, 67, 70, 99 |
| C | 3, 5, 6 | 9, 70, 73, 74, 97 |
| D | 3, 5, 6 | 99, 114, 155, 156, 159, 160 |
| E | 3, 5 | 97, 114, 147, 152, 156 |
| F | 9 | 77, 80, 81, 84, 95, 123, 143, 146, 147 |

**Figure 5:**
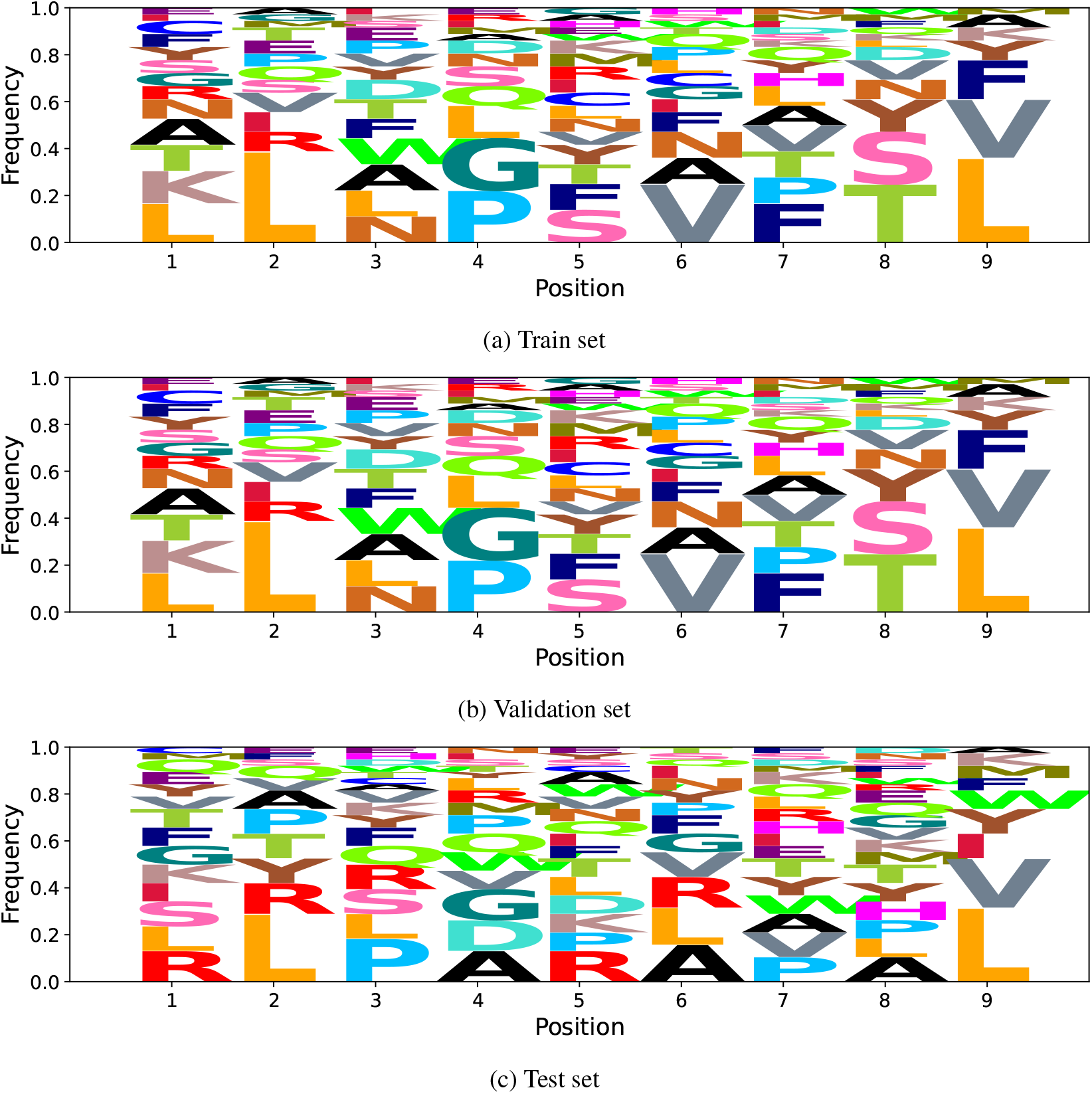
Sequence logo plots representing the amino acid frequencies at each peptide position and for each data subset.

## C Distance to Binding Pockets

Binding pockets are areas on the MHC chain that play a key role in how the peptide binds to the MHC [Nguyen et al., 2021]. Each binding pocket corresponds to a group of residues, named anchor residues that can bind to one or two peptide residues. Nguyen et al. [2021] have identified six distinct binding pockets (A to F), along with their corresponding anchor residues (on the MHC chain) and peptide residues. Specifically, for each binding pocket, there are two fixed sets of residues on peptide and MHC chain. Let 𝒫_*k*_ and ℳ_*k*_ denote the residue index sets for binding pocket *k* for peptide and MHC, respectively. The *C*_*α*_ position on these residues are therefore denoted by 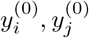 for ground truth structure and 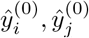 for predicted structure. In this work, we assess how well the structures reflect the position of the peptide relatively to these binding pockets by measuring the average *C*_*α*_ −*C*_*α*_ distance between the corresponding peptide and anchor residues as follows:

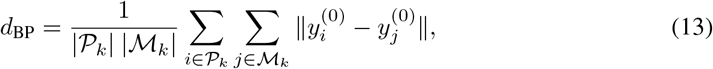

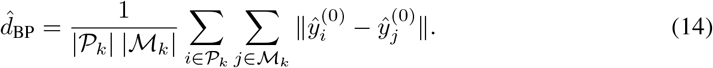

## D Experiments Parameters

GNN and training hyper-parameters are defined in Table 7 and Table 8, respectively.

**Table 7:** Neural network hyper-parameters

| Block | Number of layers | Input size | Output size |
| --- | --- | --- | --- |
| Node embedding | 1 | 15 | 128 |
| Edge embedding | 1 | 6 | 128 |
| GNN block | 4 with 4 heads each | 128 | 128 |
| Prediction head | 2 | 128/128 | 128/3 |

**Table 8:** Training hyper-parameters

| Parameter | Value | Parameter | Value |
| --- | --- | --- | --- |
| Max epochs | 70 | Output shape | (9, 5, 3) |
| Batch size | 10 | Intra-residue loss weight $\lambda_1$ | 3.0 |
| Input dimension | 15 | Inter-residue loss weight $\lambda_2$ | 0.5 |
| Embedding dimension $d_{emb}$ | 128 | Dihedral loss weight $\lambda_3$ | 0.25 |

**Table 9:** Metrics on test set per seed. The reported values are average with standard deviation across the test set.

| Metric (Unit) | Seed 1 | Seed 2 | Seed 3 | Seed 4 | Seed 5 |
| --- | --- | --- | --- | --- | --- |
| Backbone RMSD (Å) | 1.26 ± 0.64 | **1.21 ± 0.71** | 1.23 ± 0.65 | 1.23 ± 0.65 | 1.23 ± 0.62 |
| Full-atom RMSD (Å) | 2.51 ± 1.01 | **2.42 ± 1.12** | 2.43 ± 1.06 | 2.43 ± 1.06 | 2.60 ± 1.00 |
| Dihedral MAE (°) | **25.80 ± 12.55** | 30.04 ± 16.01 | 28.10 ± 13.86 | 28.10 ± 13.86 | 34.37 ± 18.10 |
| Binding Pocket A MAE (Å) | 0.11 ± 0.15 | 0.07 ± 0.14 | **0.06 ± 0.14** | 0.07 ± 0.15 | 0.06 ± 0.14 |
| Binding Pocket B MAE (Å) | 0.25 ± 0.16 | 0.20 ± 0.16 | **0.17 ± 0.15** | 0.22 ± 0.16 | 0.21 ± 0.16 |
| Binding Pocket C MAE (Å) | 0.42 ± 0.30 | 0.43 ± 0.31 | 0.39 ± 0.28 | **0.35 ± 0.29** | 0.41 ± 0.27 |
| Binding Pocket D MAE (Å) | 0.36 ± 0.36 | 0.37 ± 0.39 | **0.34 ± 0.31** | 0.37 ± 0.38 | 0.37 ± 0.34 |
| Binding Pocket E MAE (Å) | 0.49 ± 0.37 | **0.46 ± 0.32** | 0.50 ± 0.43 | 0.50 ± 0.35 | 0.54 ± 0.42 |
| Binding Pocket F MAE (Å) | 0.10 ± 0.09 | **0.09 ± 0.09** | 0.10 ± 0.09 | 0.12 ± 0.08 | 0.10 ± 0.08 |
| Attractive score MAE (kcal/mol) | **185.44 ± 83.15** | 186.44 ± 82.41 | 185.75 ± 82.92 | 185.35 ± 83.05 | 186.15 ± 83.60 |
| Total energy score MAE (kcal/mol) | 147.19 ± 53.04 | **142.43 ± 56.73** | 143.39 ± 47.87 | 143.90 ± 52.13 | 160.13 ± 82.38 |

**Table 10:** Metrics on test set per model. The reported values are average with standard deviation across the test set.

| Metric (Unit) | Full loss | MSE only | $\lambda_1 = 0$ | $\lambda_2 = 0$ | $\lambda_3 = 0$ |
| --- | --- | --- | --- | --- | --- |
| Backbone RMSD (Å) | 1.26 ± 0.64 | 1.29 ± 0.68 | 1.32 ± 0.66 | **1.24 ± 0.66** | 1.31 ± 0.71 |
| Full-atom RMSD (Å) | **2.51 ± 1.01** | 2.63 ± 1.13 | 2.67 ± 1.16 | 2.55 ± 1.14 | 2.64 ± 1.11 |
| Dihedral MAE (°) | **25.80 ± 12.55** | 31.01 ± 14.62 | 29.09 ± 16.83 | 29.01 ± 16.45 | 35.51 ± 17.57 |
| Binding Pocket A MAE (Å) | 0.11 ± 0.15 | 0.08 ± 0.15 | 0.06 ± 0.15 | **0.06 ± 0.14** | 0.07 ± 0.15 |
| Binding Pocket B MAE (Å) | 0.25 ± 0.16 | 0.29 ± 0.19 | **0.19 ± 0.15** | 0.19 ± 0.17 | 0.25 ± 0.16 |
| Binding Pocket C MAE (Å) | 0.42 ± 0.30 | 0.42 ± 0.32 | **0.41 ± 0.29** | 0.45 ± 0.34 | 0.34 ± 0.29 |
| Binding Pocket D MAE (Å) | 0.36 ± 0.36 | 0.37 ± 0.39 | 0.40 ± 0.37 | **0.35 ± 0.33** | 0.36 ± 0.35 |
| Binding Pocket E MAE (Å) | 0.49 ± 0.37 | **0.46 ± 0.38** | 0.55 ± 0.39 | 0.49 ± 0.40 | 0.47 ± 0.37 |
| Binding Pocket F MAE (Å) | **0.10 ± 0.09** | 0.11 ± 0.10 | 0.11 ± 0.09 | 0.11 ± 0.08 | **0.10 ± 0.09** |
| Attractive score MAE (kcal/mol) | **185.44 ± 83.15** | 190.20 ± 79.68 | 192.23 ± 80.19 | 190.60 ± 80.05 | 190.89 ± 80.64 |
| Total energy score MAE (kcal/mol) | **147.19 ± 53.04** | 165.00 ± 73.17 | 149.20 ± 63.33 | 168.73 ± 76.98 | 157.32 ± 57.40 |

AlphaFold 2 original and fine-tuned models were trained using Motmaen et al. [2022] implementation^1^ with original and fine-tuned weights respectively. Motmaen et al. [2022] use a 200 residue gap trick to add the peptide sequence. For OmegaFold, we use Wu et al. [2022] implementation^2^ with a 30 glycine residue gap trick as shown more successful by Tsaban et al. [2022].

## E Results

**Figure 6:**
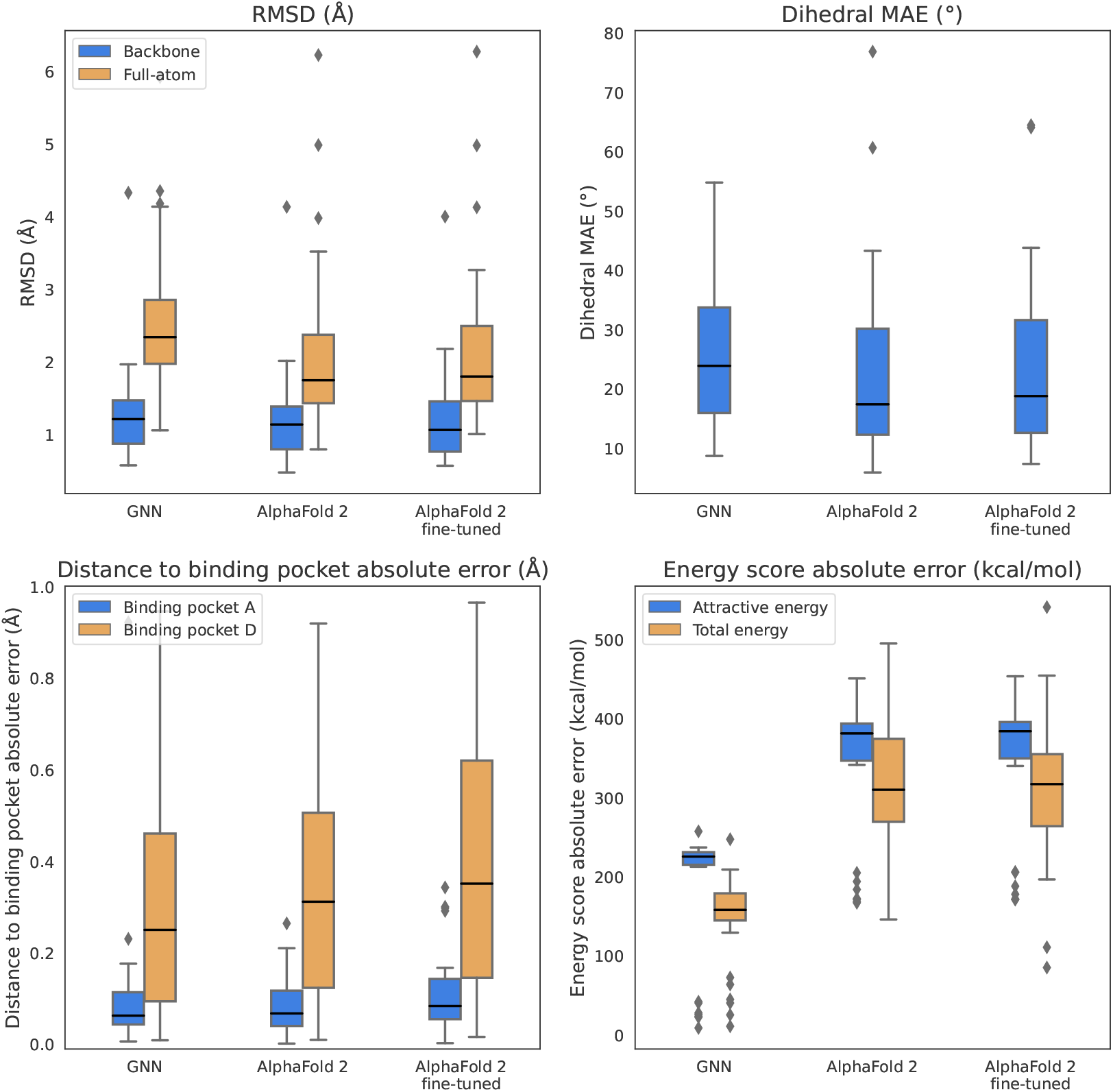
Boxplots of distributions of RMSD (top-left), dihedral MAE (top-right), distance to binding pocket absolute error (bottom-left) and energy score absolute error (bottom-right). To simplify the visualisation, OmegaFold results are not displayed and only binding pockets A and D are represented.

**Figure 7:**
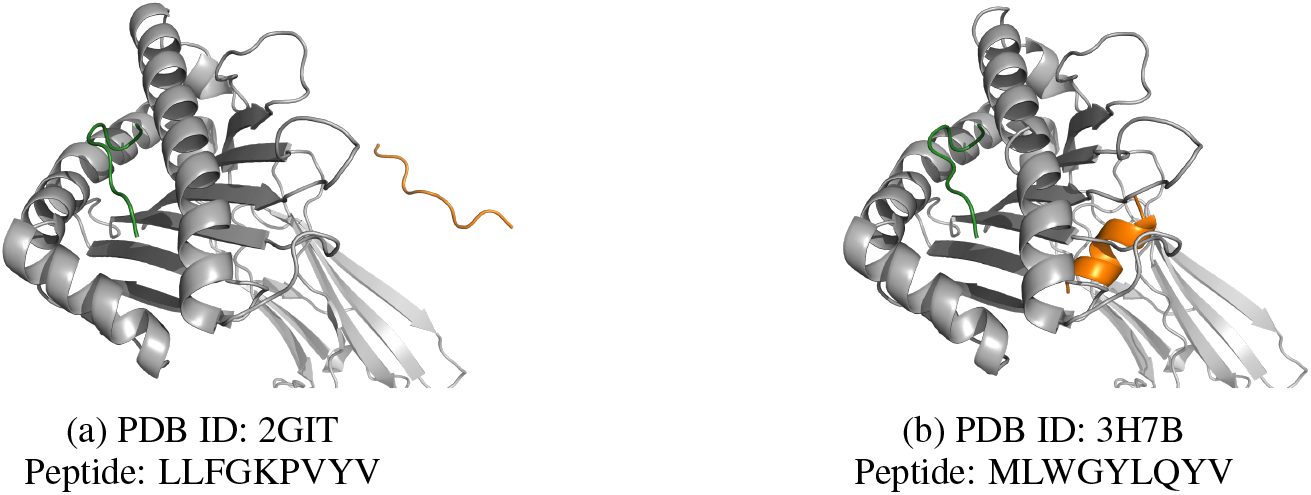
Examples of incorrectly predicted structures by OmegaFold. Experimental and OmegaFold peptide are represented in green and orange respectively. The MHC chain is represented in grey.

## F Loss Ablation

We trained the same GNN model with different combinations of loss terms. We compared different cases where additional loss terms were removed individually, and when only the MSE loss was kept. The results are reported in Table 10 and examples of predicted structures for both cases are visualised in Figure 8.

**Figure 8:**
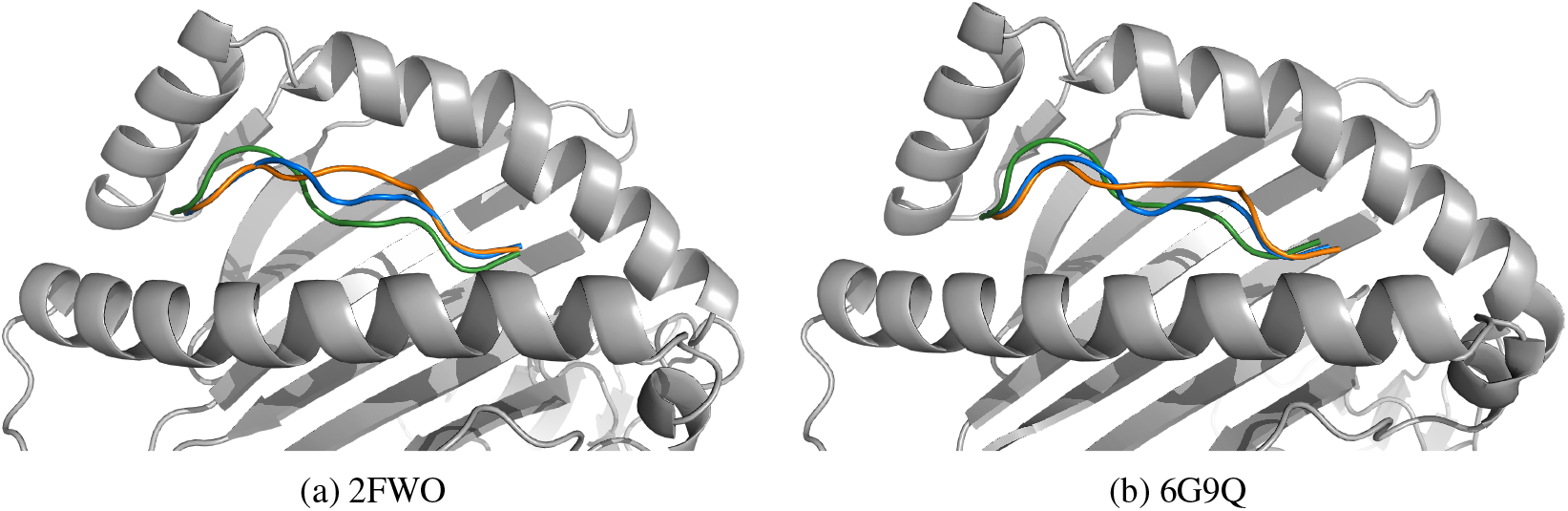
Examples of predicted structures. Experimental, GNN with full loss, and GNN with MSE loss only predicted peptide are represented in green, blue, and orange, respectively. The MHC chain is represented in grey.

## G Running time

We provide in Table 11 the inference time for each model per structure. We assess the running time on one A100 40GB GPU and 8 e2-standard 64 GB CPUs. Note that the Packer only runs on CPU and is not assessed on GPU. We hypothesise that the GNN inference running faster on CPU compared to GPU is due to the low compute requirements of the forward pass with the memory transfer from main to GPU memory dominating the cost.

**Table 11:** Inference running time in seconds per structure per model on CPU and GPU.

| Model | CPU | GPU |
| --- | --- | --- |
| GNN | 0.064 | 0.35 |
| Packer | 6.73 | NA |
| AlphaFold 2 | 303.16 | 4.61 |
| OmegaFold | 1917.00 | 26.48 |

## Footnotes

1 https://github.com/phbradley/alphafold_finetune

2 https://github.com/HeliXonProtein/OmegaFold

